# Transcriptome profiling reveals the mechanism of ripening and epidermal senescence in passion (*Passiflora edulia Sims*) fruit

**DOI:** 10.1101/2020.07.15.203968

**Authors:** Changbao Li, Ming Xin, Li Li, Xuemei He, Guomin Liu, Jiemin Li, Jinfeng Sheng, Jian Sun

**Author notes:** Corresponding authors (J. S.), (M. X.).

## Abstract

Passion fruit (*Passiflora edulia Sims*), an important tropical and sub-tropical species, is classified as a respiration climacteric fruit, the quality deteriorates rapidly after harvest. To reveal the mechanisms involved in ripening and rapidly fruit senescence, the phytochemical characteristics and RNA sequencing were conducted in the purple passion fruits with different (1-MCP and PF) treatment. Comprehensive functional annotation and KEGG enrichment analysis showed that the starch and sucrose metabolism, plant hormone signal transduction, phenylpropanoid biosynthesis, flavonid biosynthesis, carotenoid biosynthesis were involved in fruit ripening. Applying with PF and 1-MCP significantly affected transcript levels of passion fruit after harvest storage. A large number of differently expressed unigenes (DEGs) were identified significantly enrichen in starch and sucrose metabolism, plant hormone signal transduction and phenylpropanoid biosynthesis at postharvest stage. The preservative film (PF) and 1-Methylcyclopropene (1-MCP) treatments increased superoxide dismutase (SOD), catalase (CAT) and peroxidase (POD) gene expression and enzyme activities, accelerated the lignin accumulation, decline β-galactosidase (β-Gal), polygalacturonase (PG) and cellulose activities and gene expression to delay cell wall degradation during fruit senescence. The RNA sequencing data of cell wall metabolism and hormone signal transduction pathway related unigenes were verified by RT-qPCR. The results indicated that the cell wall metabolism and hormone signal pathways were notably related to passion fruit ripening. PF and 1-MCP treatment might inhibited ethylene signaling and regulated cell wall metabolism pathways to inhibited cell wall degradation. Our results reveal ripening and senescence related networks during passion fruit ripening, which can provide a foundation for understanding the molecular mechanisms underlying PF and 1-MCP treatment on fruit ripening.

## Introduction

Passion fruit (*Passiflora edulia Sims*) is an important tropical and sub-tropical species with high commercial value due to the attractive flavor and aroma, nutrients and medicinal properties for human health, such as vitamin C, B1 and B2 and essential amino acids, as well as minerals and fibers^[1, 2]^. It can be planted and grown in virtually regions and it with high demand in the market for fresh fruit and in processed foods and juices ^[3]^. The consumption of passion fruits has been reported with high contents of phytochemicals with antioxidant, anti-inflammatory and anticancer properties, which associated with lower risk of cardiovascular diseases, cancer and metabolic diseases ^[4, 5]^.

Fruit ripening and senescence, unique to plants, are complex biological processes. According to ripening and senescence processes, fruit was divided into two groups, climacteric and non-climacteric fruit ^[6]^. Passion fruit, quality deteriorates rapidly after harvest, is classified as climacteric fruit as manifested by high respiration rate, pericarp shrivel, weight loss, flavor changed and pathogens increased, which can cause tremendous economic loss and significantly restricts the supply chain of commodities ^[7]^. During fruit development, ripening is considered as a functionally modified form of senescence associated with ROS accumulation, respiration rate, ethylene production rate, and phyto-hormone level. Therefore, mature fruit with high activity of antioxidant enzymes may possibly present a prolong fruit storage life and maintenance of quality attributes for longer periods ^[8, 9]^.

In recent years, a large number of studies have been reported postharvest conservation of various fresh fruits. The method of chemical or physical preservatives were widely used to conserve fresh fruit, due to its simplest and least expensive ^[3]^. For climacteric fruit, ethylene production rate was usually increased rapidly during storage, which may accelerate the process of fruit softening or lead to physiological disorders ^[10]^. Previous studies reported that the postharvest treatment of 1-Methylcyclopropene (1-MCP) can effectively delay fruit ripening and softening process, and prolong storage life of various fruits^[11, 12]^, due to it can bind to ethylene receptors in plant cells, and inhibit ethylene action of fruit by suppressing the ethylene biosynthesis related genes expressions^[13, 14]^. The effect of 1-MCP treatment on postharvest fruits was involved in many factors, including fruit cultivars ^[15]^, concentration ^[16]^ and exposure time ^[17]^. In order to reduce the addition of chemicals on fruit or food, the preservative film (PF) provides an effective way to store fruits after harvest. However, few studies have paid attention to the postharvest storage of passion fruits by application of 1-MCP or PF. Until now, there is still no study has reported to suppress passion fruit senescence and softening during post-storage treatment of PF and 1-MCP.

Although certain studies have been reported to reduction for postharvest conservation of passion fruit by low temperature ^[3, 18]^, little concrete study available reveals the mechanism and factor-associated signaling pathways during ripening and senescence process. To better understand the mechanisms involved in the ripening and rapid fruit senescence, in present study, the phytochemical characteristics and RNA sequencing were conducted in the purple passion fruits with different (1-MCP and PF) treatment. Our results can provide reference and baseline information to assist in development of ripening and postharvest management protocols for passion fruit.

## Materials and Methods

### Plant sample collection and treatment

Passion fruits (*Passiflora edulia Sims*), ‘Tainong1’, were obtained from orchard in Guangxi Academy of Agricultural Sciences, China. The pericarp color of passion fruit was green to purple during the process of ripening. Samples were collected for different growth stage and different treatment in ripening storage, which were divided into 9 groups (A, B, C, D, E, F, G, H and J). A, B and C group were 4 weeks, 6 weeks and 8 weeks (mature stage) post anthesis (WPA), respectively. After the harvest, D and E group were stored under ambient temperature conditions (20–25 °C and 75–80% relative humidity) at 4 days and 8 days, respectively. F and G group were preservative film (PF) treated with 4 days and 8 days after ripening, respectively. For H and J group were 0.5ul/L 1-MCP treated with 4 days and 8 days after ripening, respectively. All samples stored at room temperature with 20–25 °C and 75–80% relative humidity. Three biological replicates were performed for each time points. Each peel (pericarp) of passion fruit was surface sterilized with 3% H_2_O_2_ for 10 mins, vigorously rinsed with distilled water (>200 ml/per time) for 5 times. After ground to powder in liquid nitrogen, the peel sample was stored at −80 °C for further analysis.

### Physiological index determination of passion fruit

Certain important physiological indexes of ripening and senescence in passion fruit, including the membrane permeability (Relative electrical conductivity), respiration rate and weight loss (shrinkage rate) were determined in the present study. Fifty passion fruits were selected to determine the relative electrical conductivity (REC), respiration rate and weight loss rate. The REC was measured using a conductivity meter following Wang et al.’s method ^[19]^. The respiration rate was determined according to Yang et al. ^[20]^ method. We use a TEL-7001 Infrared CO_2_ Analyzer (Telaire, Goleta, CA, USA) to determine the respiration rate of passion fruit, and expressed as mg CO_2_·kg^−1^ fresh weight (FW)·h^−1^. The weight of the each fruits was recorded on 0 d and 10 d. The weight loss rate were determined as follows: (Initial weight before storage-weight after storage)/Initial weight before storage×100%.

### Antioxidant and cell wall related enzyme activities determination

In present study, we determined the activities of antioxidant enzyme during the process of ripening and senescence in each treatment group. These antioxidant enzymes including superoxide dismutase (SOD), catalase (CAT) and peroxidase (POD) were tested in our study. The extractions and activities of SOD, CAT and POD were determined according to Chen et al. and Liu et al.’s method ^[21, 22]^ with some modification. Frozen pericarp tissue of passion fruit was homogenized add into 5 ml of 80% (v/v) ethanol, then centrifuged at 12,000×g for 10 min at 4 °C, using supernatant for antioxidant capacity determination. SOD, CAT and POD activities were measured by the absorbance at 560 nm, 240 nm and 470 nm, respectively.

The activities of β-galactosidase (β-Gal), polygalacturonase (PG) and cellulose were measured in passion fruit at different time points. The crude enzymes was extracted according to the method described by Fan et al.^[23]^ and Chen et al.^[24]^ with some modification to determine the activities of β-Gal, PG, and cellulase. The results were expressed as U/kg.

### RNA extraction and RNA_sequencing analysis

The total RNA were extracted from pericarp tissues of passion fruit using a mirVana miRNA Isolation Kit (Ambion) according to the manufacturer’s instructions. A, B, C, D, E, F, G, H and J group by control and PF or 1-MCP treated samples of passion fruit were used for RNA library preparation and sequencing. The first-strand cDNA synthesis was performed using random oligonucleotides and SuperScript II. The second-strand cDNA synthesis was performed using DNA polymerase I and ribonuclease H. Then the cDNA libraries were sequenced on an Illumina sequencing platform (HiSeqTM 2500 or Illumina HiSeq X Ten). Three biological replicates samples from each time point were used for library construction and each library was sequenced once. The raw data containing adaptors and poly-N and low-quality reads were removed. Then sequence duplication level of the clean reads were assembled into expressed sequence tag clusters (contigs) and de novo assembled into transcript and the Q20, GC content were calculated by using Trinity^[25]^. Blastx (E-value< 0.00001) was employed to search for homologues of our assembled unigenes and annotated in protein databases including NR, KOG, SwissProt and PFAM and database. The best results were used to determine the sequence orientations of the unigenes. The functional annotation by GO terms (http://www.geneontology.org) was analyzed using the program Blast2GO. The COG and KEGG pathway annotations were performed using Blastall software against the COG and KEGG databases, respectively ^[26]^.

### Real-time qRT-PCR verification

To verify the usability of the transcriptomic data, the relative transcript levels of 18 genes that were either significantly up- or down-regulated were determined using quantitative real-time reverse-transcription PCR (qRT-PCR) (TableS1). These genes included signal transduction, coloration, cell wall function, respiration and energy. Total RNA was reverse-transcribed to obtain first strand cDNA using the RevertAid First Strand cDNA Synthesis Kit (Fermentas, Lithuania) according to the manufacturer’s instructions. Gene-specific primer pairs had been designed using Primer 3.0. For qRT-PCR was performed using the SYBR^®^ Green PCR kit (Qiagen, 204054). Plant GAPDH was used as the internal reference to normalize the cDNA content. All genes were repeated at 3 times. The mRNA expression level of genes was calculated using the 2^−ΔΔCt^ method.

### Statistical analysis

Three biological replicates were used for RNA_seqencing and qRT-PCR analysis. All data for each passion fruit sample were statistically analyzed using Student’s t-test (P< 0.05). Differentially expressed unigenes were defined as unigenes with FDR<0.001 and fold change >2. P < 0.05 was considered significantly when identifying enriched GO terms and enriched KEGG pathways.

## Results

### Effect of PF and 1-MCP treatment on physiological biochemical index of passion fruit

Relative electrical conductivity (REC) in the CK, 1-MCP- and PF-treated passion fruit increased on days 0D-10D (Figure1A). However, the increase rate in REC was significantly slower in PF-treated group, compared to 1-MCP-treated at 2D-10D. Respiratory rate increased in both PF-, 1-MCP-treated and control in the passion fruit at 0D-10D. And respiratory rate of passion fruit that treated with 1-MCP and PF were significantly lower than CK group at 4D, 8D and 10D (Figure1B). The weight loss (shrinkage rate) of the control and 1-MCP-treated passion fruit showed a rapid increase from 0D to 10D, while it notably suppressed by PF treatment (Figure1C). The control group showed notably higher shrinkage rate than the 1-MCP and PF-treated passion fruit, flowing storage for 2D-10D. The results of membrane permeability, respiratory rate and weight loss in PF treated fruit were significantly lower than in 1-MCP and CK groups at 2D-10D.

**Fig 1.**
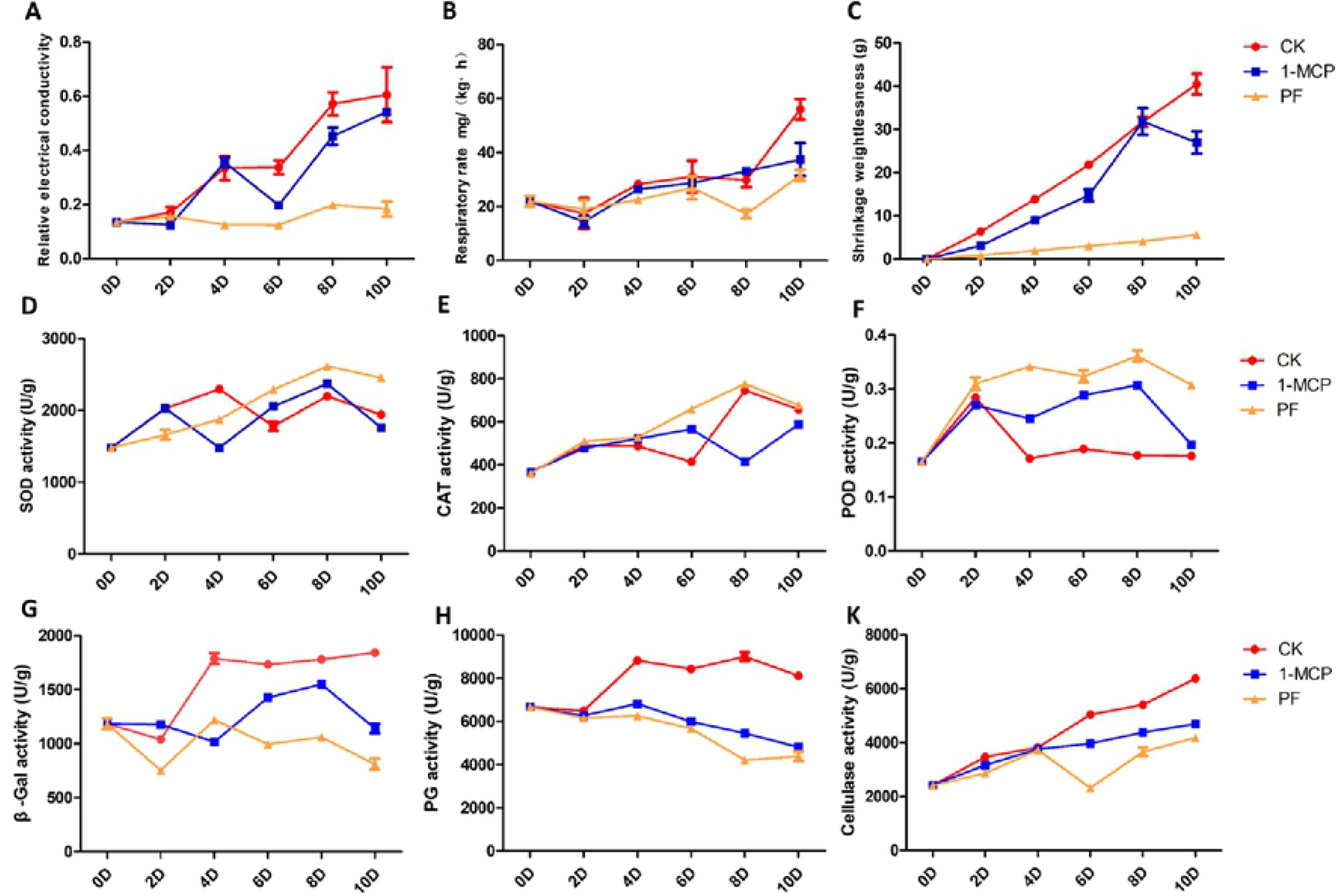

We determined ROS-scavenging enzymes activities, including SOD, POD and CAT. SOD activity increased during 0D-2D of storage, a slow decline from 4D in 1-MCP treated and 6D in CK group, then a sharp increase in the later stages at 8D and a slow decline from 10D (Figure1D). The result showed that PF treatment can activated the SOD activity, compared with the 1-MCP and CK from 6D-10D. The CAT activity was higher in PF-treated fruit than 1-MCP and control group (Figure1E). For PF treatment, the POD activity was also higher than in 1-MCP and control fruit (Figue1F).

At all the storage time points, the activity of β-Gal was significantly lower in PF-treated fruit, compared with the 1-MCP and control (Figure1G). The activity of PG was declined in both PF- and 1-MCP-treated passion fruit at 0D-10D. However, PG activity was higher in control fruit, compared with PF- and 1-MCP treated group. Cellulase activity of passion fruit was increased in both 1-MCP and PF treatment group (Figure1H). And in PF-treated fruit, cellulase activity was significantly lower than 1-MCP and control group at 4D-10D (Figure1K). The results were indicated that PF and 1-MCP treatment can significantly delay degradation of cell wall components and postpone senescence of passion fruit during postharvest storage by reduction of β-Gal, PG and cellulose activities.

### RNA-seq data analysis and function annotation of *Passiflora edulia Sims*

Transcriptome sequencing were conducted to analysis the underlying mechanisms of maturation and senescence in passion fruit by 1-MCP and PF-treatment. All stages of these samples were renamed as A (4 weeks post anthesis), B (6 weeks post anthesis), C (8 weeks post anthesis--physiological maturity), D (store at room temperature for 4 days), E (store at room temperature for 8 days), F (PF treatment for 4 days), G (PF treatment for 8 days), H (1-MCP treatment for 4 days) and J (1-MCP treatment for 8 days), respectively (Figure2). The Q30 of these samples were 92.85%- 94.65%, and 8.12 G-9.22 G raw reads were produced in different stages of passion fruit. The average GC content was 50.36% and a total of 56628 unigenes were detected in all samples after removal of low-quality reads. The average length of these unigenes was 854bp. And these unigenes were annotated in seven database. Among them 54.87% were annotated in GO, 23.01% were annotated in KO, 23.36% were annotated in KOG, 46.39% were annotated in NR, 50.37% were annotated in NT, 54.87% were annotated in PFAM and 56.74% were annotated in SwissProt (Figure3A). The results of GO and KEGG annotation were show in Figure3. We found that lipid metabolism, carbohydrate metabolism and amino acid metabolism were enriched in KEGG (Figure3B). And metabolic process, single-organism process and catalytic activity were enriched in GO (Figure3D). Expression analysis of RNA-seq data in *Passiflora edulia Sims*

**Fig 2.**
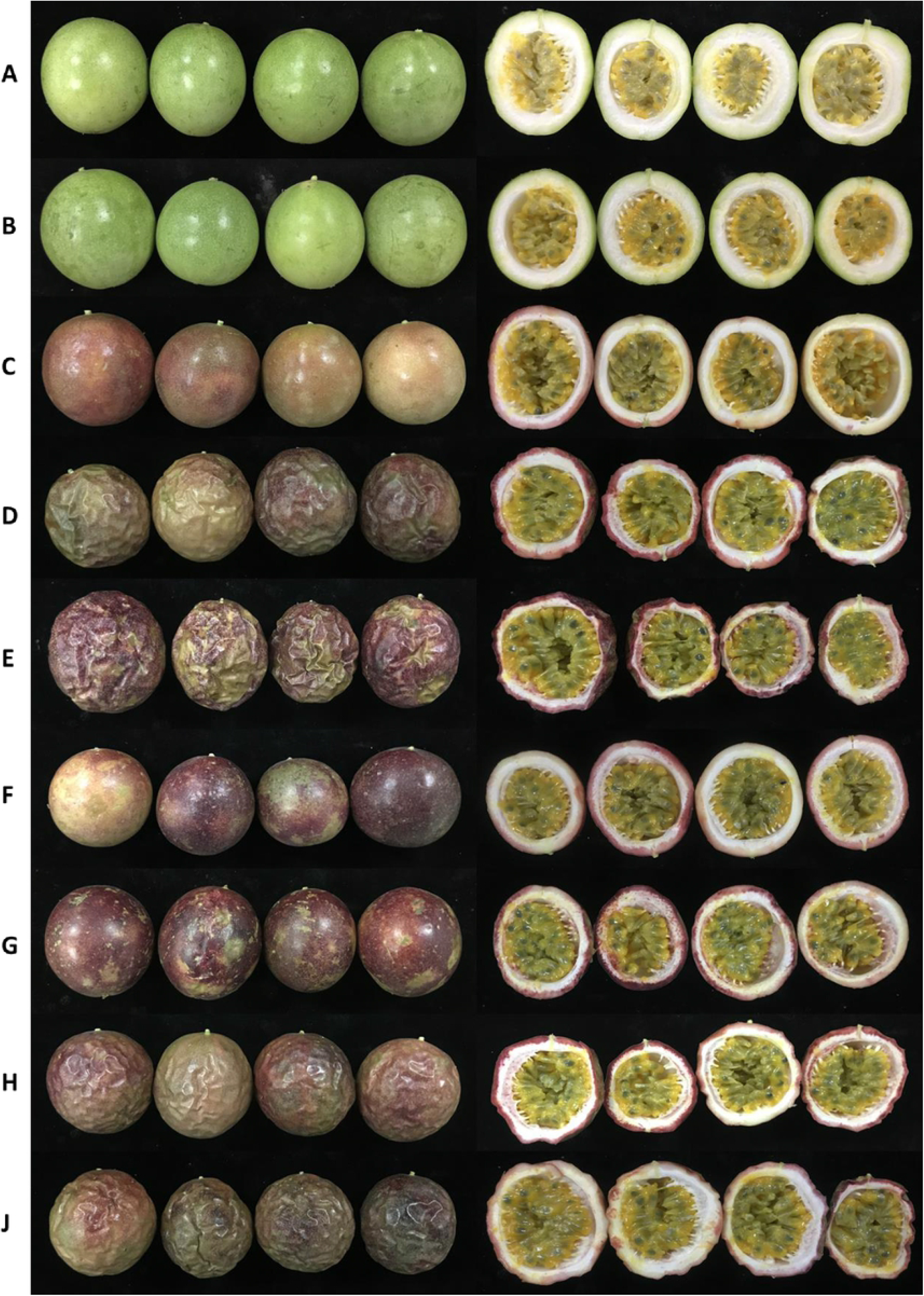

**Fig 3.**
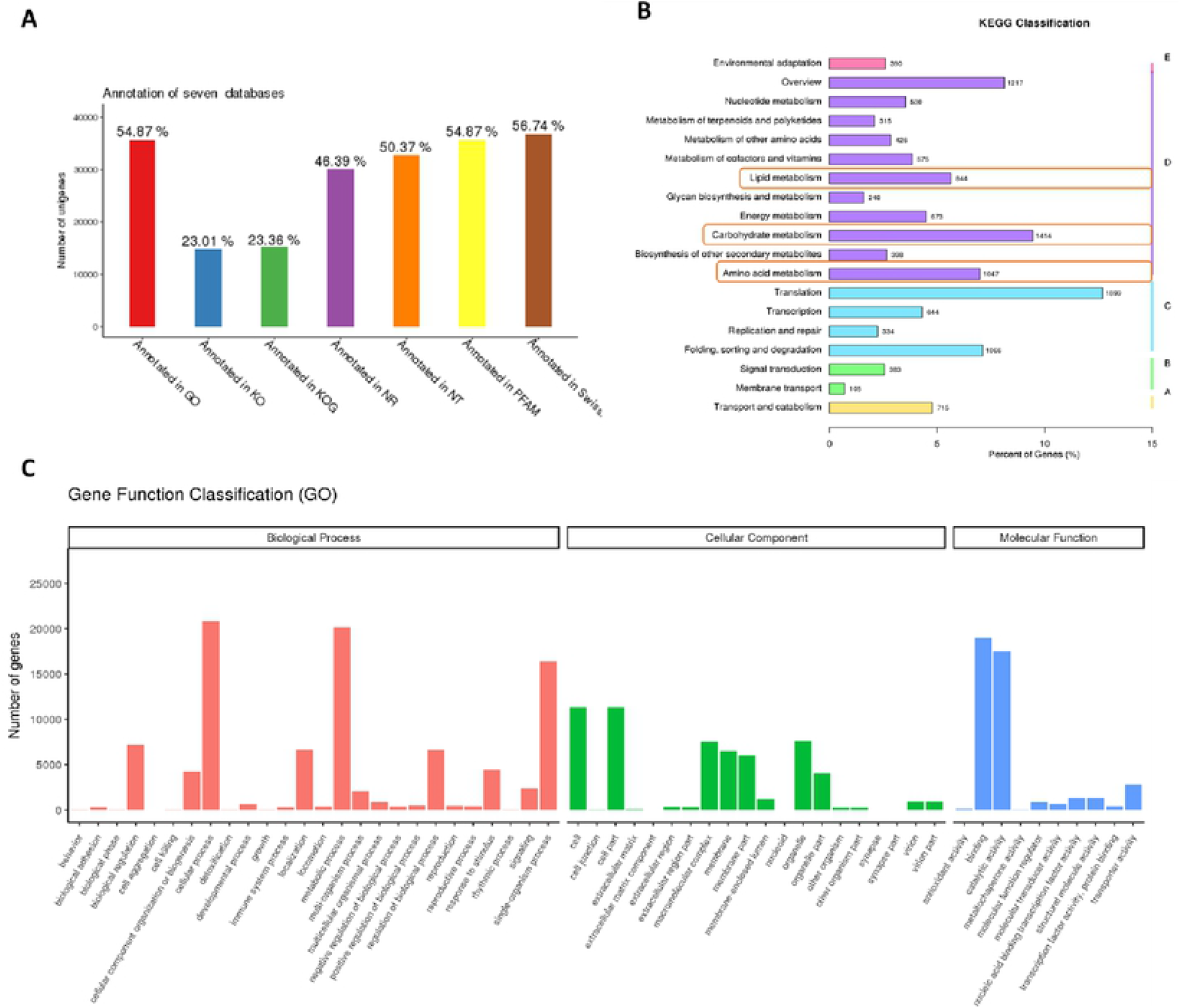

Principal component analysis (PCA) showed that the first principal component (PC1) could explain 59.36% of total variance and distinguish samples based on the time of storage at 8D (E) with other groups. E group was significantly differences with other group. At the stage of 8D, the J group (1-MCP treatment) was closely to E group, compared with G group. G group (PF treatment) was closely to C and D groups. Then the second principal component (PC2) could explain 12.4% of the total variance and separated different stages of passion fruit according to 5 times point (FigureS1A). The results suggesting that 1-MCP and PF treatment may delay the senescence of passion fruit, and the effective of PF-treated group was better than 1-MCP-treated group.

The DEGs were identified in A-J group of passion fruit with comparing the FPKMs values of each unigenes that based on the criteria log2 (Fold change) ≥2 and significance p <0.005. The result was consist with the result of PCA analysis. The expression levels of DEGs in E group (CK for 8D) was notably differentially with G (PF treatment for 8D) and J (1-MCP treatment for 8D) group. However, the expression levels of DEGs in D group (CK for 4D) was not significantly changed comparing with F (PF treatment for 4D) and H (1-MCP treatment for 4D) group (Figure S1B). Therefore, we speculated that the 8D was a key stage to study the underlying mechanism of senescence in passion fruit with PF and 1-MCP treatment.

### KEGG pathway analysis of DEGs during ripening stage

Firstly, we analysis the DEGs expression pattern during mature stage of passion fruit at A, B and C. Significant differences DEGs between B and A group and between C and A group were found for 3189 and 6622, respectively (Figure 4A). Among the 9811 DEGs, 2481 were identified in both B vs A and C vs A comparisons. Therefore, 708 and 4141 unique DEGs were expressed in C vs A and B vs A comparisons, respectively (Figure 4A). The 2481 DEGs might be involved in process of mature in passion fruit. In order to analyze the biological pathways of DEGS, we use these EDGs for Kyoto Encyclopedia of Genes and Genomes (KEGG) annotations. Then, we further use these DEGs to KEGG pathway analysis. The top 20 KEGG pathways with the most significant enrichment were shown in Figure 4B. Among all KEGG pathways, most of these DEGs were significantly enriched into ‘Starch and sucrose metabolism’ (ko00500), ‘Plant hormone signal transduction’ (ko04075), ‘Phenylpropanoid biosynthesis’ (ko00940), ‘Flavonid biosynthesis’ (ko00941), ‘Carotenoid biosynthesis’ (ko00906), ‘Biosynthesis of unsaturated fatty acids’ (ko01040), and ‘Diterpenoid biosynthesis’ (ko00904) etc. The results indicated that these pathways were mainly participated in maturation of passion fruit. With the process of passion fruit development, the color of peel were changed significantly (Figure 4) at C group. And among these pathways, the ‘Phenylpropanoid biosynthesis’ (ko00940), ‘Flavonid biosynthesis’ (ko00941) and ‘Carotenoid biosynthesis’ (ko00906) were related to fruit coloration. Therefore, we further analysis the expression level of DEGs in these two pathways to explore the potential mechanism of coloration in passion fruit during maturation.

**Fig 4.**
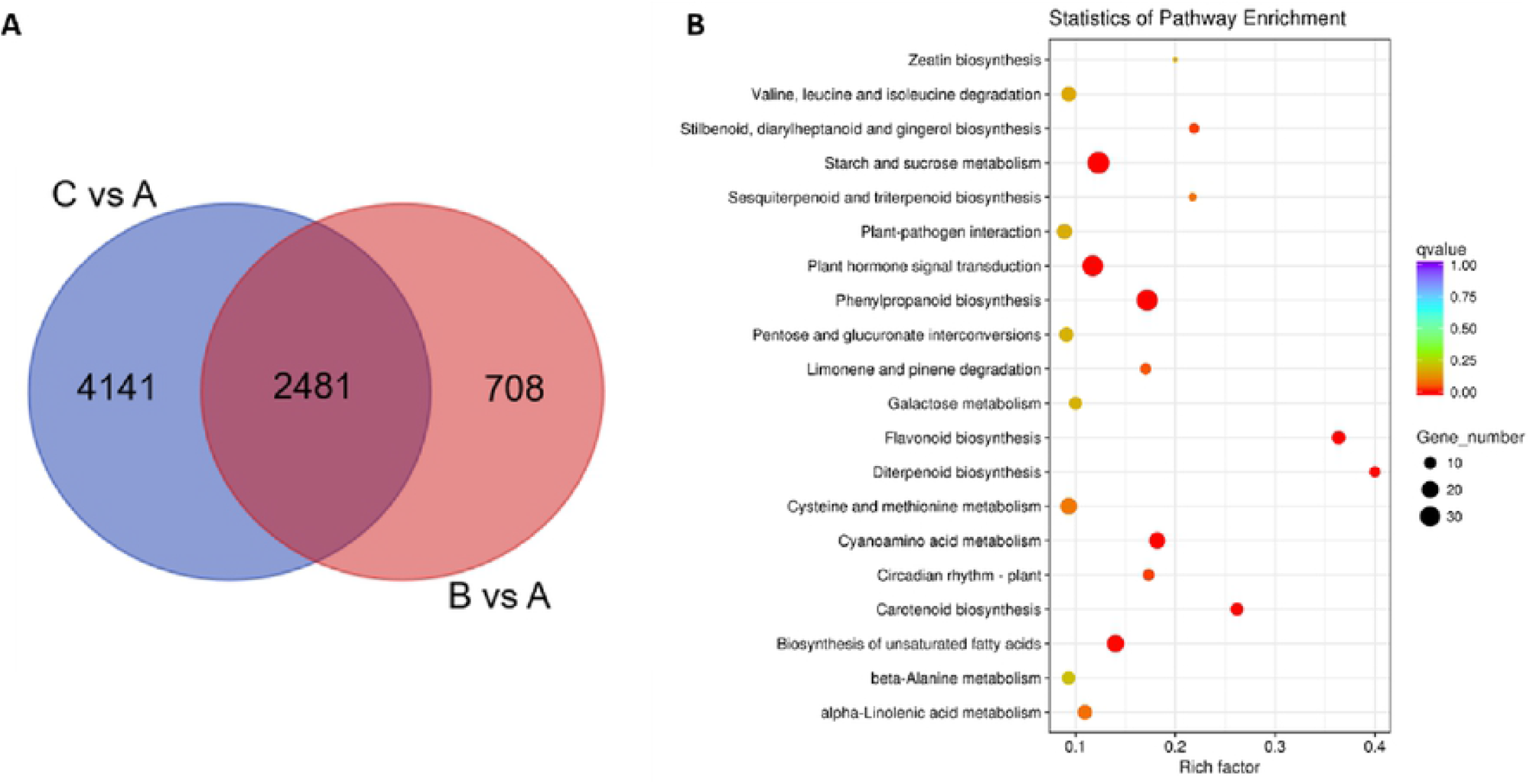

### Pericarp color changes during ripening stage

A total of 12 DEGs were identified to regulated ‘Flavonid biosynthesis’ (ko00941) in present study. Compared with A group, these 12 DEGs were up-regulated in B and C group during passion fruit development (Figure 5A, 5C), which including certain key enzyme unigenes, such as CHS (Cluster-29126.6929), ANR (Cluster-29126.6305), and DFR (Cluster-29126.9769) et al. At the same time, 11 DEGs were also identified that involved in ‘Carotenoid biosynthesis’ (ko00906), including CrtB (Cluster-29126.19339), CrtZ (Cluster-29126.16831), and NCED (Cluster-29126.17170) (Figure 5B). The expression level of these unigenes in B and C group were higher more than 10-fold comparing with A group. Therefore, the results suggesting that these unigenes may play an important roles to regulate the coloration of passion fruit during its growth and ripening. In addition, we also found that certain DEGs were significantly enriched into ‘Phenylpropanoid biosynthesis’ (ko00940) during the development of passion fruit. We all know that the downstream of this pathway was to synthesis of lignin and anthocyanin (flavonids). In this study, certain key regulated unigenes in ‘Phenylpropanoid biosynthesis’ were identified significantly up-regulated in the stage of B and C, compared with A. Among them, CCR (Cluster-29126.13688) and F5H (Cluster-29126.6512) were up-regulated significantly, which involved in the regulated degree of lignification in passion fruit during its development (Figure 5C).Thus, we inferred that the degree of lignification was also related to ripening or senescence in passion fruit.

**Fig 5.**
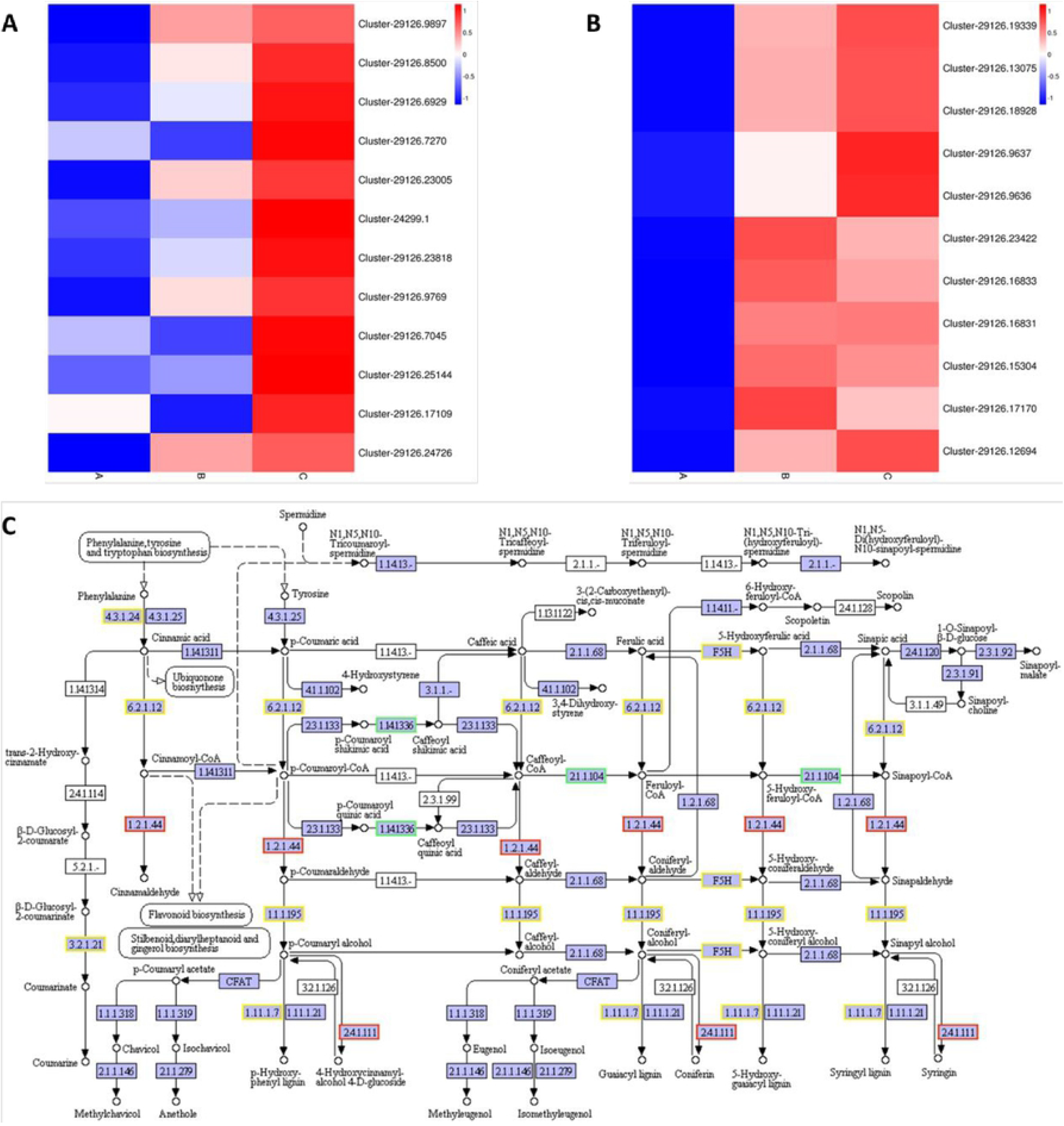

### KEGG analysis of DEGs in postharvest storage

Dynamic processes of passion fruit ripening and senescence were identified by enriched KEGG pathways of control, 1-MCP and PF treatment. For 8D of storage (control), the DEGs of E vs C significantly enriched in ‘Starch and sucrose metabolism’ (ko00500), ‘Plant hormone signal transduction’ (ko04075), ‘Phenylpropanoid biosynthesis’ (ko00940) and ‘Flavonid biosynthesis’ (ko00941). For 1-MCP and PF treated group, the DEGs of G vs E and J vs E were also significantly enriched in ‘Starch and sucrose metabolism’, ‘Plant hormone signal transduction’, ‘Phenylpropanoid biosynthesis’ and ‘Flavonid biosynthesis’ (Figure 6). These results indicated that ‘Phenylpropanoid biosynthesis’, ‘Starch and sucrose metabolism’ and ‘Plant hormone signal transduction’ were play a vital roles to regulated fruit senescence. Then in the result of KEGG pathway analysis, unigenes involved in ‘Starch and sucrose metabolism’ and ‘Plant hormone signal transduction’ were further analyzed in particular.

**Fig 6.**
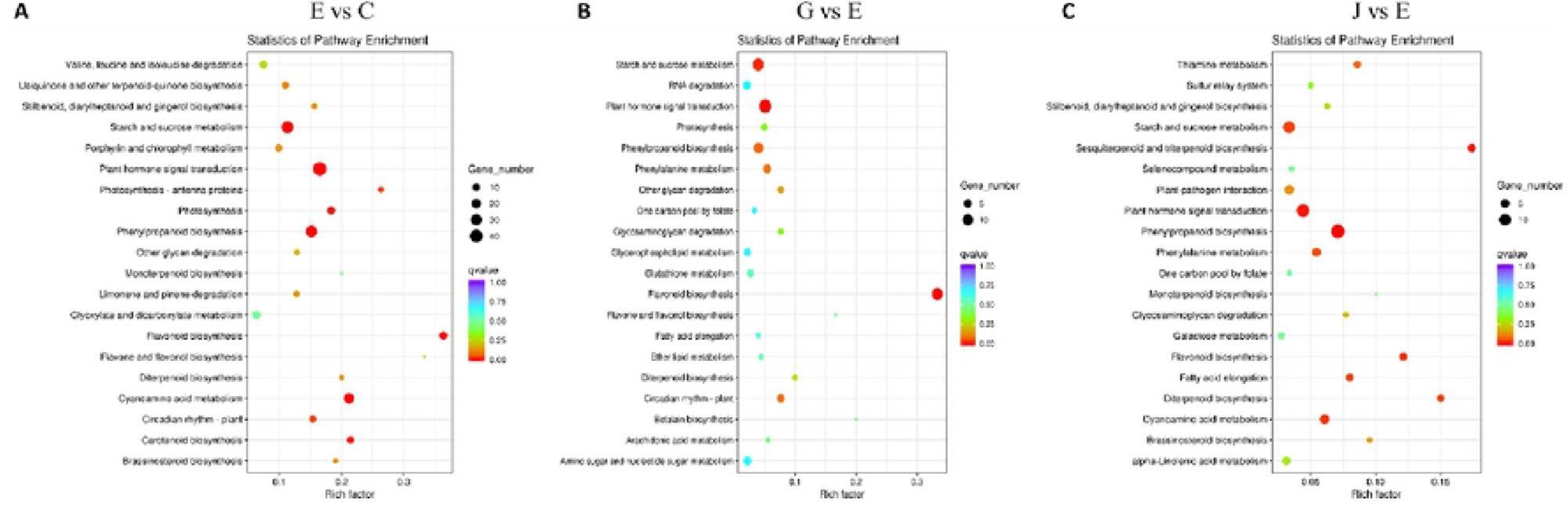

Cell wall metabolism and signal transduction pathways involved in senescence of passion fruit Most of the differentially expressed genes (DEGs) were enriched into cell wall metabolism-related pathways (Supplementary file SF2, 3, Table S1), including sucrose synthase, cellulose synthesis, pectin metabolism (PG), hemicellulose metabolism (β-Gal), and lignin metabolism (PAL, 4CL, CHS, CCR and POD). In ‘Starch and sucrose metabolism’ (ko00500) pathway, the expression levels of cluster-29126.797, cluster-29126.15857, cluster-29126.19858 and cluster-29126.23754 el al. were significantly up-regulated at process of mature stage (both B and C group). However, these DEGs were all down-regulated in G and J group by 1-MCP and PF treatment, respectively (Figure 7A). And we found that the expression level of these DEGs in G group (PF treatment for 8D) were notably higher than J group (1-MCP treatment for 8D). The DEGs in ‘Starch and sucrose metabolism’ pathway including E3.2.1.21 (beta-glucosidase), E2.4.1.14 (sucrose-phosphate synthase), E3.1.1.11 (pectinesterase) and amyA (alpha-amylase) etc.

**Fig 7.**
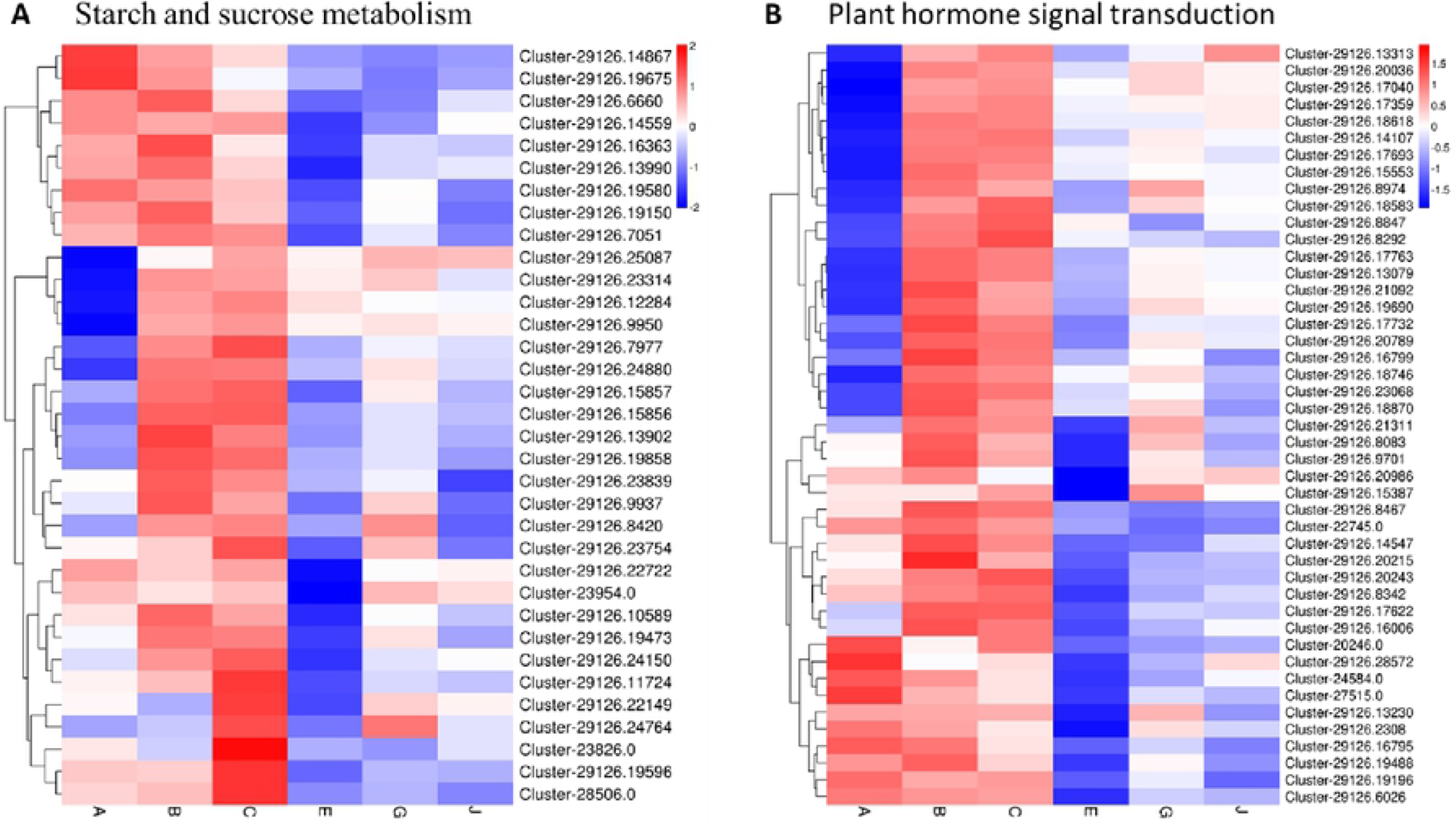

Similar with the ‘Starch and sucrose metabolism’ pathway, the expression level of DEGs were enriched in ‘Plant hormone signal transduction’ (ko04075) were analyzed in this study. A total of 45 DEGs of ‘Plant hormone signal transduction’ pathway were up-regulated during passion fruit ripening, while these DEGs were down-regulated in process of senescence (Figure 7B). And we found that the expression level of these DEGs of PF- and 1-MCP-treated (G and J) groups were up-regulated comparing with control (E) group at 8D. The DEGs were key unigenes to regulated ethylene and auxin synthesis, which include certain key enzyme unigenes, such as ETR, EBF1_2, and CTR1 etc. These results indicated that the cell wall metabolism and plant hormone signal transduction (ethylene and auxin) pathway were essential to regulated passion fruit ripening and senescence. And we also found that PF and 1-MCP treatment can affected the expression of these cell wall metabolism related unigenes, to delay the senescence of passion fruit.

### Transcriptome validation through qRT-PCR

To validate the transcriptome sequencing data, eighteen unigenes with different expression patterns were selected for qRT-PCR analysis (Figure S2). These genes included three ‘Carotenoid biosynthesis’ related genes (Cluster-29126.19339, Cluster-29126.16831, Cluster-29126.17170); four ‘Flavonid biosynthesis’ related genes (Cluster-29126.6929, Cluster-29126.23818, Cluster-29126.9769, Cluster-29126.22876); three ‘Phenylpropanoid biosynthesis’ related genes (Cluster-29126.13688, Cluster-29126.5571, Cluster-29126.6512); three ‘Plant hormone signal transduction’ related genes (Cluster-29126.17622, Cluster-29126.21896, Cluster-29126.8974); two ‘Starch and sucrose metabolism’ related genes (Cluster-29126.10589, Cluster-29126.16363); three ‘Cell wall synthesis and degradation’ related genes (Cluster-29126.23818, Cluster-29126.22988, Cluster-29126.10249). These genes are involved in signal transduction, coloration, cell wall function, respiration and energy, and other types of metabolism. The results of RNA sequencing and qRT-PCR data showed a positive correlation in this study (R^2^=0.564, p<0.01, Figure S2). It indicated that the RNA_Seq data was reliable to further analysis.

## Discussion

### Changes of pericarp color and related molecular mechanisms during ripening

The pericarp color changed from green to purple in passion fruit during ripening stage. In stage of A, B and C of passion fruit, the expression pattern of DEGs were significantly enriched into ‘Phenylpropanoid biosynthesis’, ‘Flavonid biosynthesis and ‘Carotenoid biosynthesis’. A large number of study reported that ‘Flavonid biosynthesis’, ‘Carotenoid biosynthesis’, and downstream of the ‘Phenylpropanoid biosynthesis’ pathway were involved in coloration of fruit^[27, 28]^, by anthocyanin and carotenoid synthesis and accumulation. The content of anthocyanin and carotenoid contribute to the color of the fruit that can contribute to human health-promoting properties.

### Changes of physiological index and related enzymes at postharvest storage

Climacteric fruit usually suffers from rapid senescence after harvest, which exhibits increase in weight loss, respiratory rate, ethylene production rate and physiological disorders at room temperature storage. The results will lead to fruit quality deterioration, tissue softening, dehydration and flavor changing ^[29, 30]^. Purple passion fruit exhibited a typical climacteric pattern during postharvest ripening ^[31]^. Therefore, in order to study the mechanism of fruit senescence and preservation during storage process, we determined the physiological and biochemical indexes in passion fruit by PF and 1-MCP treatment at room temperature storage. Firstly, weight loss and respiratory rate were determined in different storage time points with PF or 1-MCP treatment. Respiratory burst was elicited through aerobic and anaerobic respiration, which was regulated specifically by related signal transduction. In this study, weight loss and respiratory rate increased progressively and peaked at the later stage of storage at room temperature. However, this increased trend was significantly inhibited by 1-MCP and PF treatment, and lowest weight loss and respiratory rate were shown in PF-treated fruit. The respiratory burst was mainly related to increased production of reactive oxygen species (ROS), to enhanced peroxidase activity and stimulating lipoxygenase pathway ^[32]^, which were closely related to accelerated senescence of fruit.

ROS accumulation level was controlled by the balance between capacity of ROS production and scavenging ^[33]^. The cell membrane, maintain relative stability in the internal environment, might be damaged by excess ROS level which also cause peroxidization and accelerates senescence in fruit ^[34]^. Therefore, the enzymatic and non-enzymatic ROS scavenging systems to scavenge the potential damage to the cells ^[35, 36]^. The antioxidant enzymes of ROS scavenger including superoxide dismutase (SOD), peroxidase (POD), and catalase (CAT) etc ^[37]^. Our result have revealed that applying 1-MCP and PF postharvest could enhance SOD, POD, and CAT activities in passion fruit, and the related gene expression pattern were also consist with RNA_seq analysis. In addition, fruit also contains a variety of antioxidants, including phenolic compounds and anthocyanins ^[38]^.

### Plant hormones involved in fruit senescence at postharvest storage

Besides ROS, other signaling pathways or molecular are also involved in plant senescence, including plant hormones transduction, ethylene, auxin, jasmonic acid and salicylic acid ^[39]^. Plant hormones are indispensable to regulate fruit ripening and senescence, which controlled fruit color, sugar, flavor and aroma during ripening and senescence ^[40]^. And previous study reported that ethylene and auxin were plays a major role in the ripening and senescence process of climacteric fruits ^[41]^. In present study, the result revealed that a large number of DEGs were enriched in plant hormone signal transduction, and up-regulated in ethylene and auxin synthesis pathways during fruit ripening (B and C), while significantly down-regulated at senescence process at postharvest storage. These DEGs including certain ethylene and auxin synthesis regulated key receptor unigenes ETR, EBF1_2, and CTR1 etc. PF and 1-MCP treatment can mitigate the low expression trend of these DEGs in passion fruit at postharvest storage.

### Membrane and cell wall components changes at postharvest storage

The process of fruit senescence after harvest will leading to the irreversible destruction of membrane integrity and to accelerated leakage of ions ^[42]^. Loss of membrane integrity may lead to subcellular de-compartmentalization which resulting in enzymatic browning catalyzed by peroxidase and polyphenol oxidase in postharvest fruits ^[43]^. The membrane integrity damage is reflected by membrane permeability, which can be determined by the relative electrical conductivity (REC) ^[44]^. In the present study, REC increased under 1-MCP and control conditions after storage, while PF treatment notably repressed the increases REC parameters and regulating gene expression of ‘Biosynthesis of unsaturated fatty acids’ pathway. The result suggesting that the prevention of passion fruit by PF treatment might be involved in decline the oxidative destruction of membranes. Consist with our results, previous study reported that melatonin inhibited membrane phospholipid degradation and maintained the degree of unsaturation fatty acids, which contribute to the preservation of membrane integrity in tomato fruit ^[45]^. The degradation of unsaturated fatty acids results in destruction of the cell membrane integrity of the fruit peel ^[46, 47]^. PF and 1-MCP treatment inhibit membrane lipid peroxidation that was beneficial for maintaining unsaturated degradation to saturated fatty acids, which can maintain passion fruit cell membrane integrity.

Cell wall degradation is essential for fruit quality during the fruit ripening and the senescence process ^[40]^. The thickness and strength of cell wall are key components to maintenance the fruit firmness ^[48]^. In the process of fruit senescence, the progressive disassembly of the primary cell wall structure and components were depolymerized by the action of cell wall hydrolases ^[49]^. Senescence involved degradation of cell wall composition enzymes mainly include β-galactosidase (β-Gal), polygalacturonase (PG), pectin methylesterase (PME), pectin lyase (PL), cellulase and expansin ^[50, 51]^. In present study, certain cell-wall metabolism related pathways were identified by RNA_seq data, such as cellulose synthesis, pectin metabolism, and hemicellulose metabolism and most of the DEGs were significantly up-regulated during fruit ripening. Similar to RNA_seq data, the activities of β-Gal, PG and cellulase were also significantly increased in passion fruit during storage, which were all inhibited by the PF and 1-MCP treatment. Under the condition of PF treatment, the activity of these three enzymes was the lowest than 1-MCP and control group. PG was regarded as pectin-degrading enzymes in fruits ^[52]^, which leading to cell wall loosing and fruit softening ^[53]^. β-Gal was a pectin-debranching enzyme which capable to simultaneously modify pectin and hemicellulose[54]. And cellulase was widely regarded to cause the degradation of cellulose matrix in the cell walls of fruits ^[55]^. In the present results, the integrity of the cell wall was well maintained by PF and 1-MCP treatment to inhibit the fruit softening and PF-treated showed better results than 1-MCP. Therefore, our results suggesting that PF treatment can alleviate passion fruit quality deterioration and suppress the enzyme activities of β-Gal, PG and cellulase to inhibit cell wall components degradation.

### The expression of sucrose and cell wall degradation related DEGs during postharvest storage

The content of sugar, fructose and glucose are the key factors in the formation of fruit quality, which were significantly accumulated during the fruit ripening process ^[56, 57]^. However, the sucrose and fructose content of passion fruit decreased during postharvest storage ^[7]^. In this study, the DEG expression level of starch and sucrose metabolism pathway was significantly up-regulated at ripening stage in passion fruit, while these DEGs were all down-regulated in G and J group by 1-MCP and PF treatment. Therefore, we speculate that PF and 1-MCP can delay the senescence of passion fruit.

Starch, hemicelluloses, cellulose, and pectin are the major factors of cell wall polysaccharides ^[58]^. In present study, these related pathways were identified by RNA_seq, including sucrose metabolism and lignin metabolism. Most of these unigenes were involved in the cell wall polysaccharides that were significantly affected by PF and 1-MCP treatment. Lignin is one of the most abundant polyphenolic polymers in higher plants that function to the structural support of the cell walls, water tightness, and response to environmental stimulate ^[59]^. Certain studies has reported that the activities of POD, PAL, C4H, and 4CL were positively correlated with lignin accumulation in loquat fruit ^[60]^. In the present study, the lignin metabolism-related unigenes in ‘Phenylpropanoid biosynthesis’ pathway were identified by RNA_seq analysis, such as POD, PAL, C4H, and 4CL. The results has consist with the reported that several postharvest approaches, such as 1-MCP and PF could inhibit lignification in fruit ^[61]^.

In general, these results consist with previous studies that exogenous 1-MCP, melatonin or glycine betaine can delays postharvest senescence in fruits ^[47, 62]^ by reducing respiratory rates, REC, cell wall degradation key enzyme activity and increasing antioxidant enzyme activity and related gene expression, such as the activity of β-Gal, PG, cellulase, SOD, POD and CAT. The PF treatment can slows the respiratory rate of passion fruit and consequently lowers the deterioration of organoleptic traits, such as flavor, aroma and cell wall among other quality characteristics.

## Supporting information

**Figure S1.** Clustering analysis of RNA-seq samples and the intersection of differentially expressed unigenes (DEGs) among fruit ripening and postharvest stages. (A) Principal component analysis of RNA-seq profile for passion fruit developmental stages, and postharvest stages with different treatment. (B) The DEGs counts of different comparison groups, grey indicates up-regulated and blue represents down-regulated. (DOCX)

**Figure S2.** Correlation between qRT-PCR and RNA_seq for the 18 unigenes. Each point represents a fold change value of expression level between the corresponding FPKM and RT-PCR in passion fruit. (DOCX)

**Table S1.** List of primers used by RT-PCR and RT-PCR and FPKM by RNA_seq data. (XLSX)

**Supplementary File SF1.** Expression profile of DEGs expression related to flavonoids, anthocyanin and carotenoids synthesis. (XLSX)

**Supplementary File SF2.** Expression profile of DEGs expression related to phenylpropanoid biosynthesis pathway. (XLSX)

**Supplementary File SF3.** DEGs expression profile related to starch and sucrose metabolism. (XLSX)

**Supplementary File SF4.** DEGs expression profile related to plant hormone signal transduction. (XLSX)

## Authors’ contributions

CBL, MX and JS provided experimental design and plant material. CBL performed experiments and RNA_Seq data analyses. CBL and MX were identification phenotype of passion fruit and data analysis. CBL wrote the manuscript. LL, XMH, GML, JML and JFS were read this article and modified. All authors read and approved the final manuscript.

